# Targeted BDNF upregulation via upstream open reading frame disruption

**DOI:** 10.1101/2025.03.04.641482

**Authors:** Ning Feng, Thomas Goedert, Nenad Svrzikapa, Dongnan Yan, Britt Hanson, Alicia Ljungdahl, Ruxandra Dafinca, Kevin Talbot, Stephan J. Sanders, Dhanu Gupta, Mathew J.A. Wood, Thomas C. Roberts

**Affiliations:** Institute of Developmental and Regenerative Medicine, University of Oxford, IMS-Tetsuya Nakamura Building, Old Road Campus, Roosevelt Dr, Headington, Oxford, OX3 7TY, UK; Department of Paediatrics, University of Oxford, South Parks Road, Oxford, OX1 3QX, UK; Orfonyx Bio Ltd, BioEscalator, Innovation Building, Rm 10.15, University of Oxford, Roosevelt Drive, Oxford, OX3 7FZ, UK; Nuffield Department of Women’s and Reproductive Health, University of Oxford, John Radcliffe Hospital, Oxford, OX3 9DU, UK; Department of Physiology, Anatomy and Genetics, University of Oxford, South Parks Road, Oxford, OX1 3QX, UK; Oxford Motor Neuron Disease Centre, Nuffield Department of Clinical Neurosciences, University of Oxford, John Radcliffe Hospital, Oxford OX3 9DU, UK; Department of Psychiatry and Behavioral Sciences, UCSF Weill Institute for Neurosciences, University of California, San Francisco, San Francisco, CA 94158, USA; New York Genome Center, New York, NY 10013, USA; MDUK Oxford Neuromuscular Centre, UK, OX3 7TY, UK

**Author notes:** To whom correspondence should be addressed. **Corresponding Author** Dr Thomas C. Roberts, Institute of Developmental and Regenerative Medicine, University of Oxford, IMS-Tetsuya Nakamura Building, Old Road Campus, Roosevelt Dr Headington, Oxford OX3 7TY, United Kingdom. **Author Contact Information** Ning Feng, Thomas Goedert, Nenad Svrzikapa Dongnan Yan Britt Hanson Alicia Ljungdahl Ruxandra Dafinca Kevin Talbot Dhanu Gupta Stephan Sanders Matthew J. A. Wood Thomas C. Roberts.

**Keywords:** uORF, upstream open reading frame, BDNF, brain-derived neurotrophic factor, base editing

## Abstract

To understand the relative contributions of 5ʹ UTR elements to translation, we performed a comprehensive analysis of upstream open reading frames (uORFs) across a representative 5ʹ UTR. We selected the neurotrophin BDNF (Brain derived neurotrophic factor) as an exemplar as upregulation of this protein is a potential therapeutic approach for a plethora of neurodevelopmental, neurodegenerative, and neuropsychiatric disorder indications. Predicted uORFs were identified in 14 out of 17 *BDNF* RefSeq transcript isoforms, and experimentally confirmed to be exerting translation repression effects for five of these transcripts. These findings suggest that uORF elements play an important role in shaping the protein output from this locus. We explored several approaches to disrupt *BDNF* uORF function. Deletion of a 5ʹ UTR exon in *BDNF* v11 (containing eight predicted uORFs), in order to simulate an exon skipping outcome, resulted in pronounced upregulation in a reporter construct system. This effect was found to be partially uORF-dependent, but was also dependent on the disruption of an RNA secondary structure element. However, this transcript variant was found to not be expressed in human brain. Conversely, direct disruption of a single uORF start codon in the widely expressed *BDNF* v4 transcript variant using an adenine base editing approach resulted in a ∼1.8-fold upregulation of endogenous BDNF protein expression in cell culture. This study describes novel *BDNF* regulatory mechanisms, and potential uORF-targeted modalities for therapeutic gene activation.

## Introduction

Brain derived neurotrophic factor (BDNF) is a secreted growth factor of the neurotrophin family that plays an important role in both the development and maintenance of the nervous system. BDNF promotes neuronal cell survival and differentiation by binding to neurotrophic tyrosine kinase receptor 2 (NTRK2), also known as tropomyosin-related kinase B (TrK B).(1, 2) As such, BDNF contributes to learning and memory by increasing dendritic branching, long-term potentiation, and synaptic plasticity.(3–6) A common valine to methionine substitution at BDNF residue 66 (rs6265) has been reported to result in reduced BDNF protein secretion and memory deficits,(7) and reduced BDNF levels have been associated with a multitude of neurological and psychiatric disorders including Alzheimer’s disease,(8, 9) Parkinson’s disease,(10, 11) Huntington’s disease,(12–14) Rett syndrome,(15) and depression.(16–19) Upregulating BDNF levels has been shown to improve symptoms in a mouse model of Rett syndrome,(20) and reduce neuronal loss and improve pathology in various animal models of Alzheimer’s disease,(21) Parkinson’s disease,(22, 23) Huntington’s disease,(24) and amyotrophic lateral sclerosis (ALS).(25) Moreover, given its neuroprotective effects, the administration of BDNF is being investigated as a possible therapeutic strategy for the treatment of acute injuries, with the aim of limiting the extent of damage following stroke,(26, 27) spinal cord injury,(28) and traumatic brain injury.(29)

A variety of methodologies have been explored to increase BDNF levels including delivery of recombinant protein,(30) peptide mimetics of BDNF,(31) gene therapy,(32) unsilencing via targeting of a natural antisense transcript at the *BDNF* locus,(33) and small molecule repurposing.(34, 35) Indeed, recombinant BDNF protein has been investigated in clinical trials for ALS,(36) and a *BDNF* gene therapy Phase I clinical trial is currently underway for the treatment of Alzheimer’s disease (NCT05040217).

Human transcripts frequently contain upstream open reading frames (uORFs) which consist of an AUG start codon located within the 5ʹ UTR followed by an in-frame stop codon. The presence of one or more uORFs is typically associated with translational repression of the downstream primary open reading frame (pORF),(37) and, in some cases, nonsense-mediated decay.(38) Predicted uORFs are present in more than 50% of human transcripts, suggesting that a large proportion of the human transcriptome may be held in a semi-repressed state. As such, disruption of uORF-mediated regulation provides an opportunity to relieve this repression, and thereby achieve targeted upregulation of a specific therapeutically relevant protein, such as BDNF.

Here we have identified multiple uORF-regulated *BDNF* transcript isoforms, offering potential for therapeutic manipulation. uORF-mediated regulation was experimentally confirmed for five *BDNF* transcripts. *BDNF* transcript v11 (NM_001143811.2) includes a spliced 5ʹ UTR, whereby the removal of one exon was found to result in pronounced upregulation that was dependent on uORF activity and the presence of an RNA secondary structure element. Characterisation of the *BDNF* transcriptional landscape in human brain tissue identified *BDNF* transcript v4 (NM_001709.5) as the optimal target transcript, which contains two uORFs. Disruption of the start codon for one of the uORFs using an adenine base editing approach resulted in up to a 2-fold increase in endogenous BDNF protein expression. These studies describe novel regulatory mechanisms at the *BDNF* locus with potential for therapeutic exploitation. Furthermore, the approaches described here provide blueprints for targeted protein upregulation at other gene loci.

## Results

### Analysis of the transcriptional landscape and predicted uORFs at the *BDNF* locus

The human *BDNF* locus was analysed to identify predicted uORFs with publicly available ribosome profiling (Ribo-Seq)/RNA-Seq data overlaid (**Figure 1**). The *BDNF* gene consists of seventeen RefSeq transcripts, of which 14 were predicted to contain at least one uORF (**Table 1**). Thirteen of these transcripts encode an identical 248 amino acid protein, while the remaining four transcripts encode various N-terminally-extended BDNF protein isoforms. The transcripts primarily differ in their transcriptional start sites and splicing of 5ʹ UTR exons.

**Figure 1.**
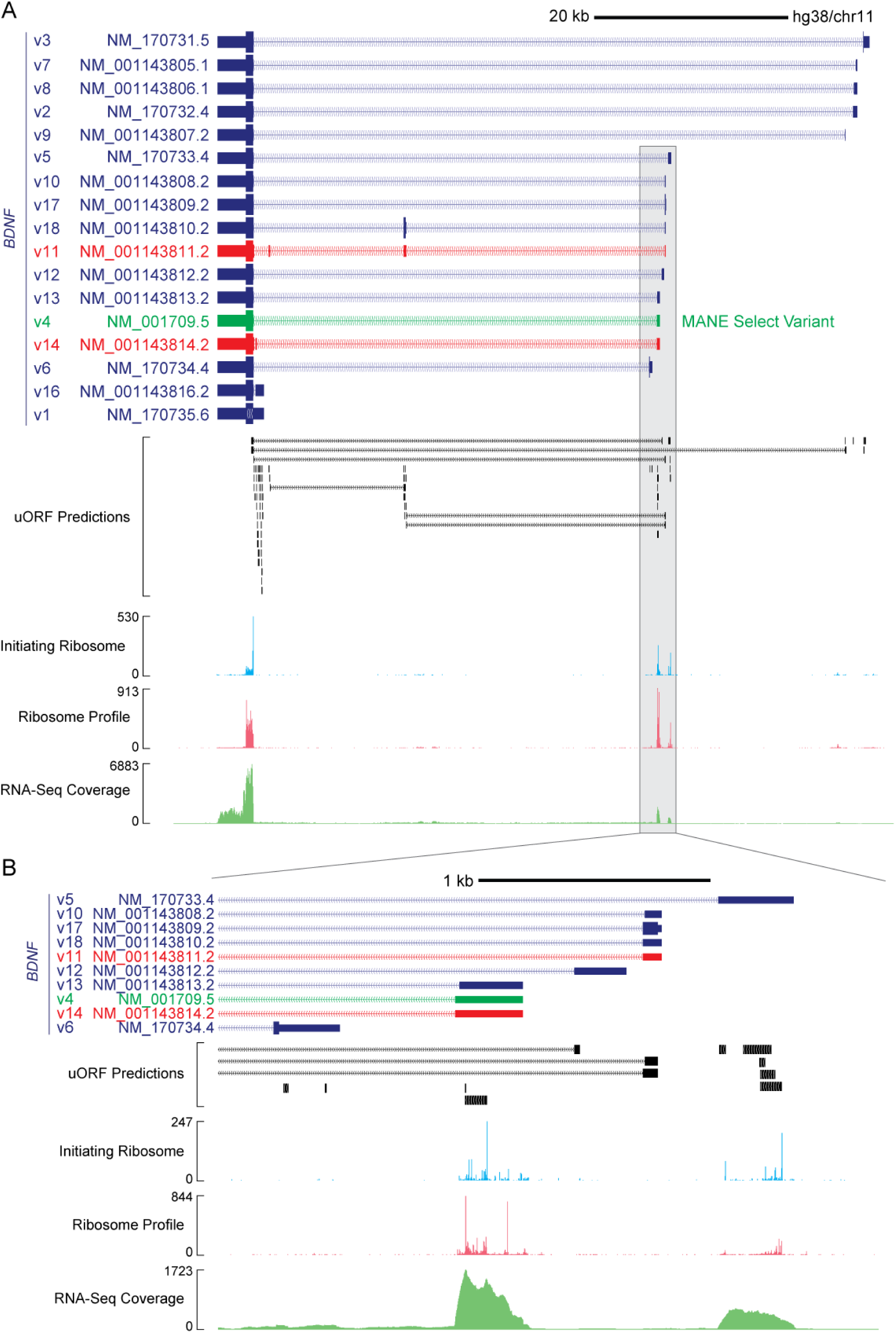
Analysis of predicted uORFs at the *BDNF* locus. (**A**) Genome browser screenshot of the *BDNF* locus showing the 17 RefSeq transcript isoforms. The MANE Select Variant is highlighted in green. Two transcripts with skippable 5ʹ UTR exons are highlighted in red (NM_001143811.2 and NM_001143814.2). Positions of predicted uORFs are indicated and the data are combined with publicly available Ribo-Seq and RNA-Seq data from the GWIPS-viz browser. (**B**) Zoomed in view of the first exons for 9 transcript isoforms with Ribo-Seq evidence of translation at certain uORFs.

**Table 1.** *BDNF* RefSeq transcript variants. Transcript variants investigated in this study are highlighted in bold.

| RefSeq ID | Transcript Variant | Other Names | Encoded protein isoform | Predicted uORFs | 5' UTR exons | uORF/exon structure | CDS size (aa) |
| --- | --- | --- | --- | --- | --- | --- | --- |
| NM_170735.6 | v1 | IX | a | 22 | 1 | [22] | 248 |
| **NM_170732.4** | **v2** | **IIc / BDNF2C** | **a** | **1** | **2** | **[1,0]** | **248** |
| NM_170731.5 | v3 | I / BDNF1 | b | 3 | 1 | [3] | 256 |
| **NM_001709.5** | **v4** | **VIb / BDNF5** | **a** | **2** | **2** | **[2,0]** | **248** |
| **NM_170733.4** | **v5** | **IV / BDNF4** | **a** | **5** | **2** | **[5,0]** | **248** |
| NM_170734.4 | v6 | VIIb / BDNF6B | c | 2 | 1 | [2] | 263 |
| NM_001143805.1 | v7 | IIa / BDNF2A | a | 0 | 2 | [0,0] | 248 |
| NM_001143806.1 | v8 | IIb / BDNF2B | a | 0 | 2 | [0,0] | 248 |
| NM_001143807.2 | v9 | III / BDNF3 | a | 1 | 2 | [1,0] | 248 |
| NM_001143808.2 | v10 | Va | a | 1 | 2 | [1,0] | 248 |
| **NM_001143811.2** | **v11** | **V-VIII-VIIIh** | **a** | **11** | **4** | **[1,8,2,0]** | **248** |
| NM_001143812.2 | v12 | Vh | a | 1 | 2 | [1,0] | 248 |
| NM_001143813.2 | v13 | VIa | a | 2 | 2 | [2,0] | 248 |
| **NM_001143814.2** | **v14** | **VIb-IXbd** | **a** | **3** | **3** | **[2,1,0]** | **248** |
| NM_001143816.2 | v16 | IXabd | a | 17 | 2 | [17,0] | 248 |
| NM_001143809.2 | v17 | Vb | d | 0 | 1 | [0] | 277 |
| NM_001143810.2 | v18 | V-VIII | e | 2 | 2 | [1,1] | 330 |

### Multiple *BDNF* transcript isoforms are regulated by uORFs

We next sought to experimentally validate uORF functionality in various *BDNF* isoforms using a dual luciferase reporter system. The appropriate *BDNF* 5ʹ UTR was cloned upstream of a Renilla luciferase transgene and mutants were generated in which uORFs were disrupted by converting their respective ATG start codons to TTG, thereby ablating their activity. A separate firefly luciferase cassette served as an internal control. Initially, we focused on the MANE Select (Matched Annotation from NCBI and EMBL-EBI) transcript isoform (*BDNF* v4, NM_001709.5, ENST00000356660.9). This transcript was predicted to contain two uORFs, the second of which is a minimal uORF (i.e. consisting of a start codon followed immediately by a stop codon) (**Figure 2A**).

**Figure 2.**
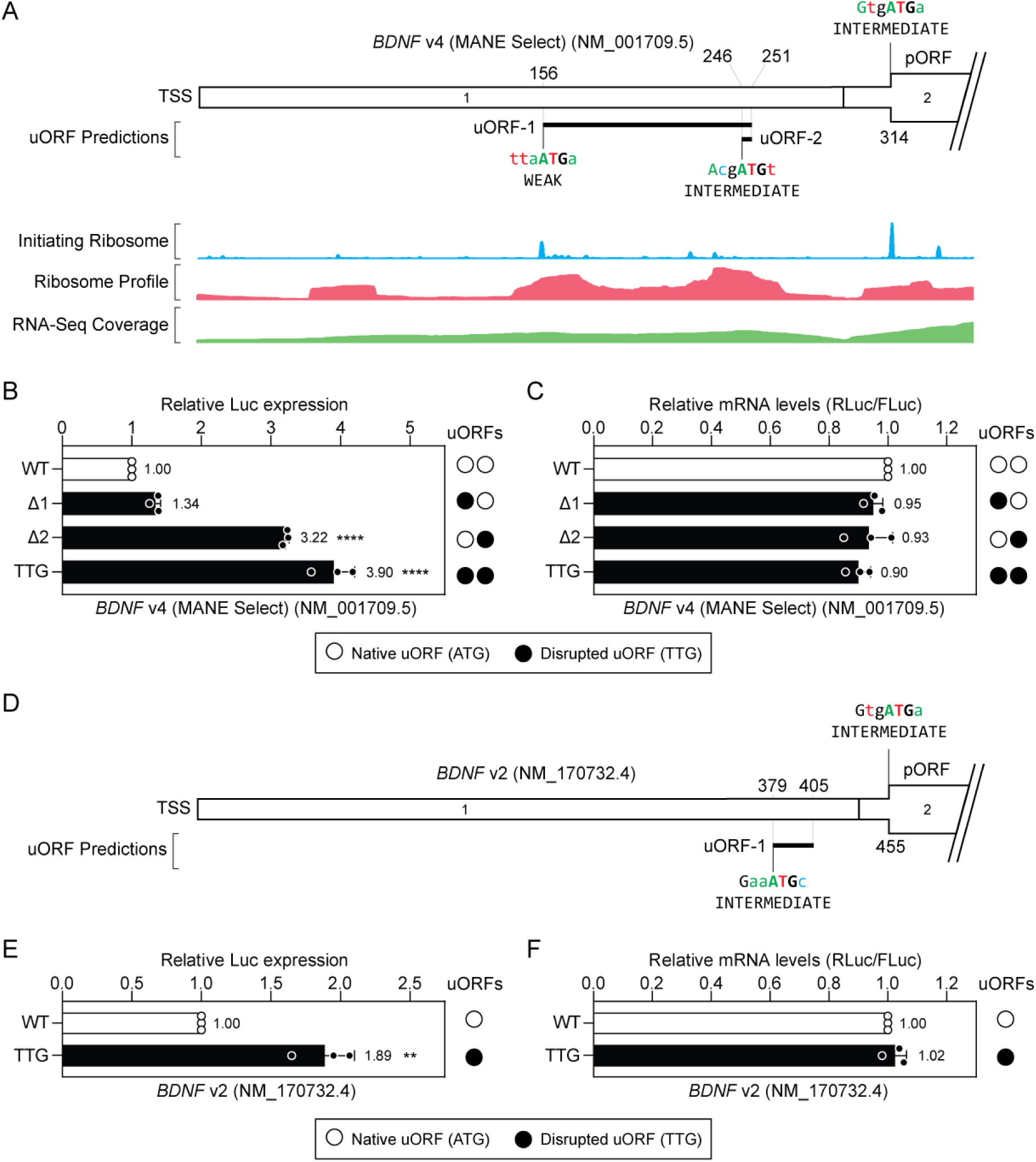
BDNF transcript isoforms are subject to uORF-mediated translational repression. (**A**) Schematic of *BDNF* transcript v4 NM_001709.5 (the MANE Select variant), containing two predicted uORFs (i.e. uORF-1 and uORF-2). Aggregated Ribo-Seq and RNA-Seq data are overlaid providing evidence of uORF translation. (**B**) HEK293T cells were transfected with *BDNF* v4 5ʹ UTR dual luciferase reporter constructs as indicated and luciferase activity assayed after 24 hours. uORFs were disrupted by mutation of the corresponding upstream ATG to TTG. (**C**) mRNA transcript levels (Renilla luciferase and firefly luciferase) were analysed in parallel by RT-qPCR. (**D**) Schematic of *BDNF* transcript v2 NM_170732.4, containing one predicted uORF (uORF-1). (**E**) HEK293T cells were transfected with *BDNF* v2 5ʹ UTR dual luciferase reporter constructs and luciferase activity assayed after 24 hours as described above. (**F**) mRNA transcript levels were determined in parallel as described above. Values are mean+SD (*n*=3 independent experiments), and were scaled such that the mean of the WT control group was returned to a value of 1. Statistical significance was determined by one-way ANOVA with Bonferroni *post hoc* test and Student’s *t*-test, as appropriate. \*\**P*<0.01, \*\*\*\**P*<0.0001.

HEK293T cells were transfected with the *BDNF* v4 reporter plasmid constructs and dual luciferase activity was assessed 24 hours later. Disruption of both uORFs resulted in a 3.9-fold increase in reporter expression (*P*<0.0001) (**Figure 2B**). Disruption of uORF-1 resulted in minimal upregulation that did not reach statistical significance, suggesting that its contribution to this repressive effect is minimal. By contrast, disruption of uORF-2 resulted in a ∼3.2-fold increase in reporter expression (*P*<0.0001). Differences in luciferase activity between experimental groups could not be explained by altered mRNA transcript levels (**Figure 2C**). These findings suggest that the *BDNF* MANE Select transcript isoform is subject to a high level of uORF-mediated translational repression. This activity is mostly due to uORF-2 (a minimal uORF), although disruption of uORF-2 alone did not reach the same level of de-repression as simultaneous disruption of both uORFs, suggesting that there is an additive repressive effect of these two uORFs in series.

A second *BDNF* transcript isoform (*BDNF* v2, NM_170732.4) was analysed in parallel. This transcript contains a single-predicted uORF in a pORF-proximal position (**Figure 2D**). Disruption of this uORF resulted in a ∼1.9-fold increase in reporter expression (*P*<0.01) (**Figure 2E**), which could not be explained by changes in mRNA transcript levels (**Figure 2F**). Both *BDNF* transcript variants v2 and v4 might potentially be targeted for therapeutic BDNF protein elevation via disruption of a single uORF.

A third *BDNF* transcript isoform (*BDNF* v5, NM_170733.4) was similarly investigated. This transcript contains five predicted uORFs (**Figure 3A**). Mutant constructs were generated in which various combinations of uORFs were disrupted as described above. Disruption of all five uORFs resulted in a ∼3.6-fold increase in reporter expression (*P*<0.0001). However, disruption of any single uORF alone resulted in minimal (or negligible) de-repression that was not significantly different from the wild-type control (**Figure 3B**). These data suggest that there is redundancy in function between uORFs arranged in series within the same 5ʹ UTR, such that this transcript is not a suitable target for a single uORF-targeted disruption approach.

**Figure 3.**
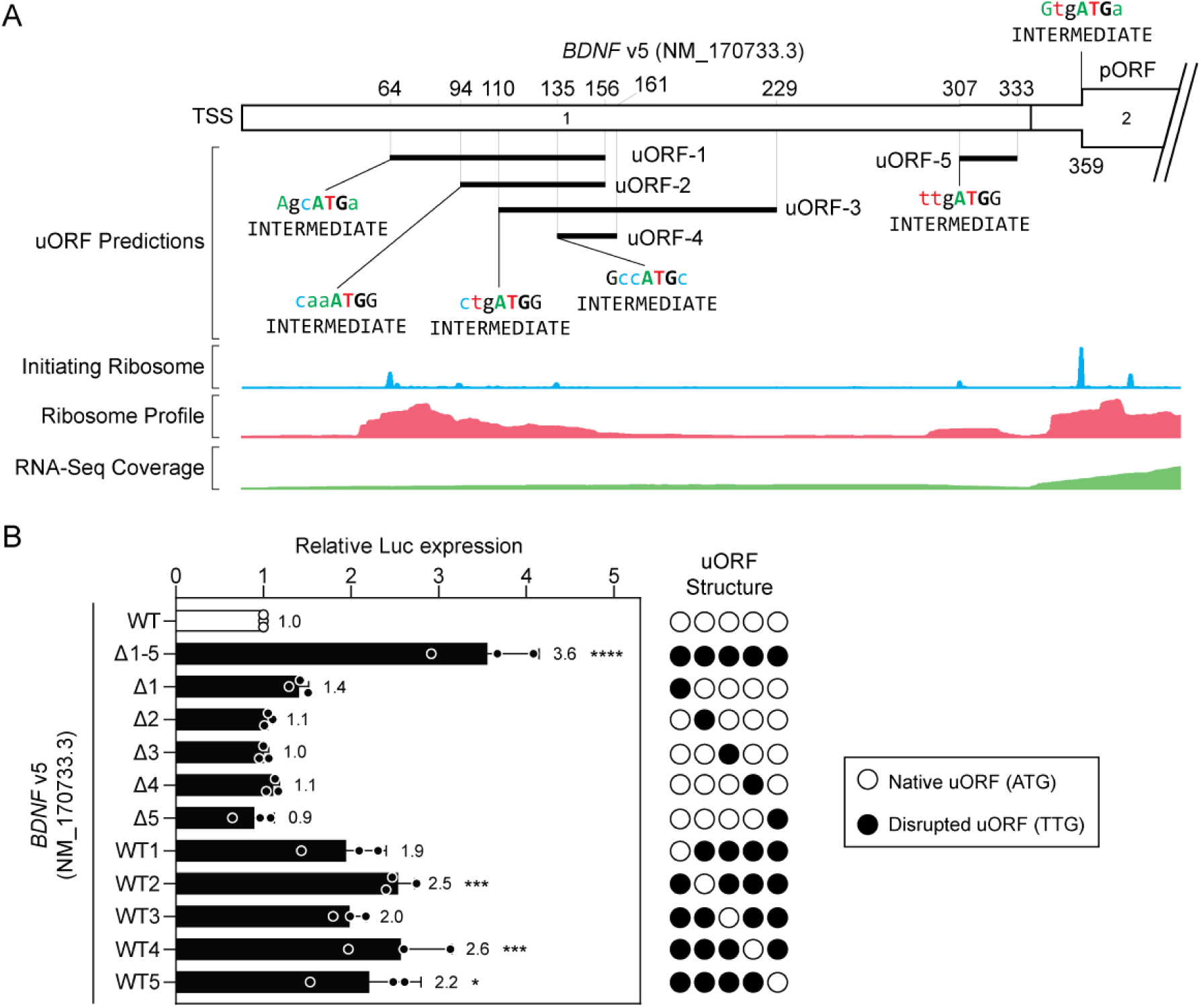
BDNF transcript v5 is subjected to uORF-mediated repression. (**A**) Schematic of *BDNF* transcript v5 NM_170733.4, containing five predicted uORFs. Aggregated Ribo-Seq and RNA-Seq data are overlaid providing evidence of uORF translation. (**B**) HEK293T cells were transfected with *BDNF* v5 5ʹ UTR dual luciferase reporter constructs as indicated and luciferase activity assayed after 24 hours. uORFs were disrupted by mutation of the corresponding upstream ATG to TTG. Values are mean+SD (*n*=3 independent experiments), and were scaled such that the mean of the WT control group was returned to a value of 1. Statistical significance was determined by one-way ANOVA with Bonferroni *post hoc* test, \**P*<0.05, \*\*\**P*<0.001, \*\*\*\**P*<0.0001.

A fourth *BDNF* transcript isoform (*BDNF* v14, NM_001143814.2) was also studied. This transcript contains the same first exon as the MANE Select isoform (*BDNF* v4), containing 2 uORFs, described above, but also contains an alternatively spliced second exon containing a third uORF (**Figure 4A**). Disruption of all three uORFs resulted in a ∼2.9-fold increase in reporter expression (*P*<0.001), whereas disruption of uORF-2 resulted in a ∼2.1-fold increase (*P*<0.05) (**Figure 4B**). Conversely, a construct containing only uORF-2 was found to be highly repressive, leading to reporter levels that were not significantly different from the WT control construct. These data suggest that uORF-2 is responsible for the majority of translation repression activity. Notably, uORF-2 is the same minimal uORF (i.e. a start codon immediately followed by a stop codon) that was found to be repressive in the context of the MANE Select variant (*BDNF* v4) described above (**Figure 2B**).

**Figure 4.**
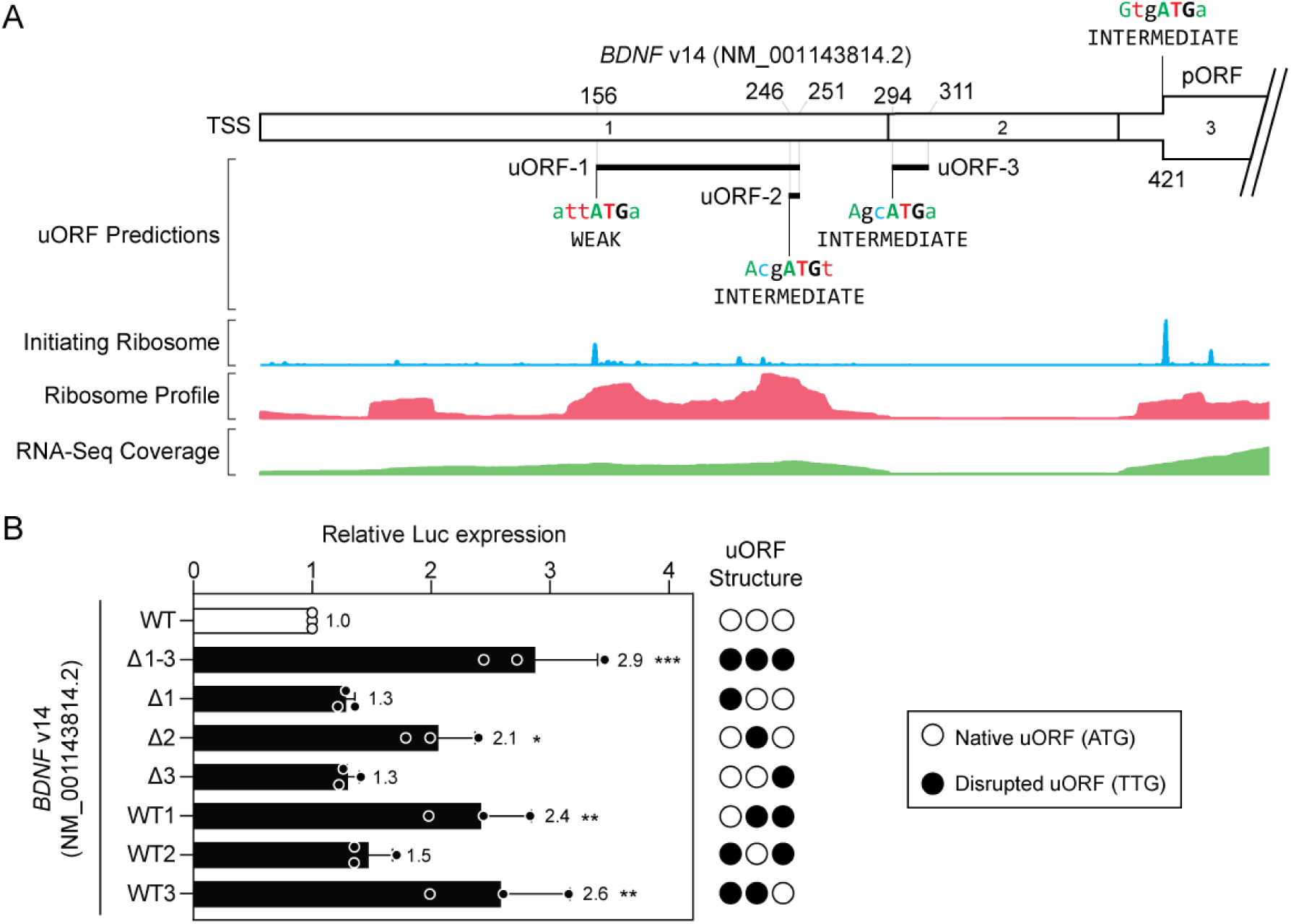
BDNF transcript v14 is subjected to uORF-mediated repression. (**A**) Schematic of *BDNF* transcript v14 NM_001143814.2, containing three predicted uORFs. Aggregated Ribo-Seq and RNA-Seq data are overlaid providing evidence of uORF translation. (**B**) HEK293T cells were transfected with *BDNF* v14 5ʹ UTR dual luciferase reporter constructs as indicated and luciferase activity assayed after 24 hours. uORFs were disrupted by mutation of the corresponding upstream ATG to TTG. Values are mean+SD (*n*=3 independent experiments), and were scaled such that the mean of the WT control group was returned to a value of 1. Statistical significance was determined by one-way ANOVA with Bonferroni *post hoc* test, \**P*<0.05, \*\**P*<0.01, \*\*\**P*<0.001.

### Potential for BDNF upregulation via upstream exon exclusion

Splice modulating oligonucleotides targeting the 5ʹ UTR have the potential to exclude untranslated exons containing uORFs from the resulting mature mRNA transcript, potentially resulting in therapeutic de-repression of the downstream pORF. As such, two *BDNF* transcripts were identified that qualified as putative exon skipping targets: transcript v11 (NM_001143811.2) and transcript v14 (NM_001143814.2). These transcripts contained at least three 5ʹ UTR exons, with one or more ‘skippable’ exons. In this context, skippable exons are defined as 5ʹ UTR exons that do not contain the transcription start site or pORF translation initiation site, and which contain at least one uORF initiation codon.

The 5ʹ UTRs of each of these transcripts were cloned downstream of a Renilla luciferase gene as described above, and variants generated whereby each skippable exon, or combination of exons, was deleted. All constructs were transfected in HEK293T cells and luciferase activity measured 24 hours post transfection.

For *BDNF* v11, deletion of exon 2 (containing eight predicted uORFs) resulted in a ∼9.8-fold upregulation of pORF reporter gene expression (*P*<0.01) (**Figure 5A,B**). Deletion of both exons 2 and 3 resulted in even further de-repression, with a ∼19.1-fold increase in Renilla luciferase activity (*P*<0.0001). However, deletion of exon 3 alone (containing two predicted uORFs) did not affect pORF reporter gene expression. The profound upregulation in luciferase activity observed could not be explained by changes in transcript levels as there was no difference observed between groups (**Figure 5C**).

**Figure 5.**
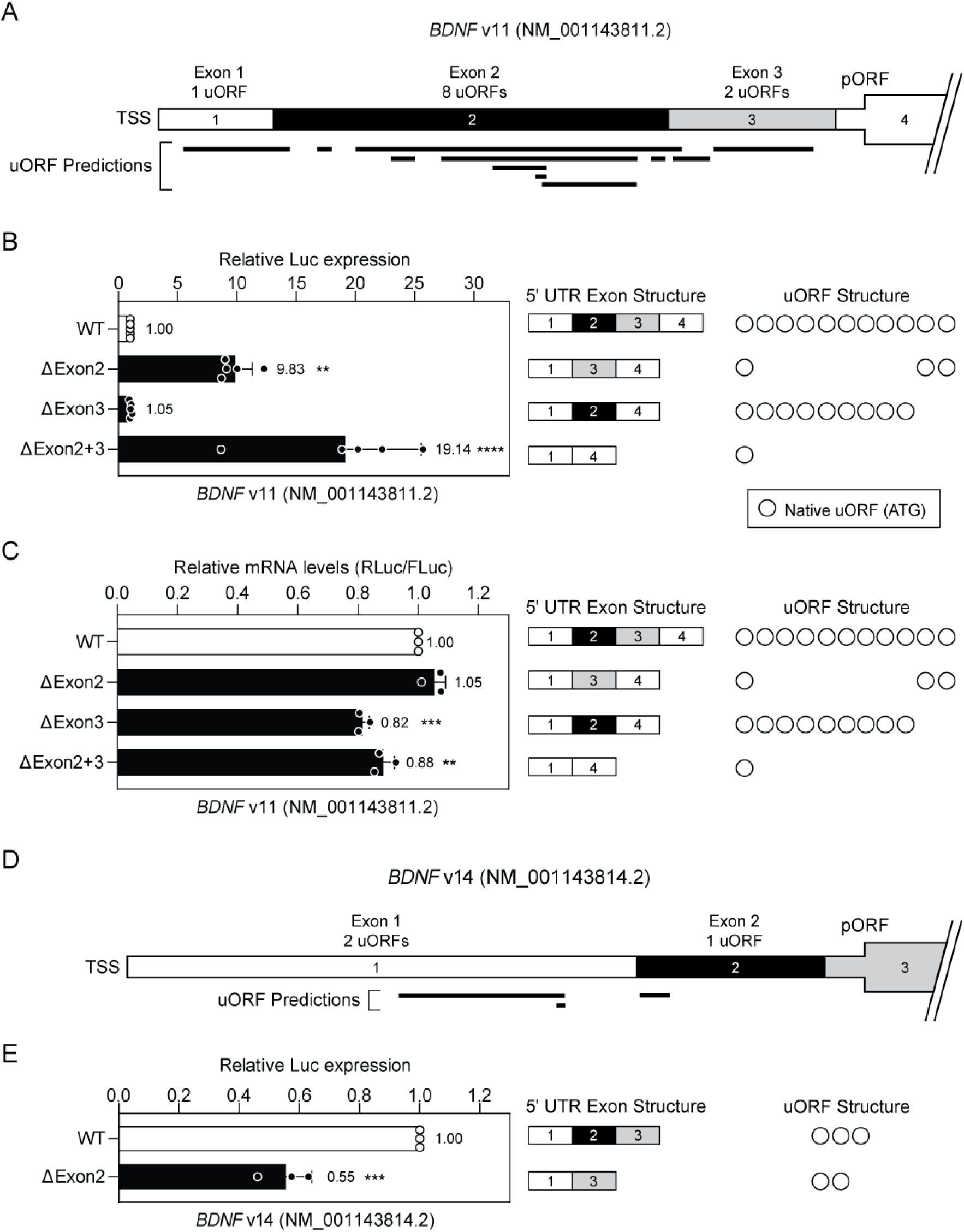
Effect of exon deletion on *BDNF* primary ORF translation. (**A**) Schematic of the 5ʹ UTR for the *BDNF* v11 transcript (NM_001143811.2) indicating the sizes and positions of exons, the locations of predicted uORFs, and the number of uORFs per exon. The pORF start codon is located in exon 4. (**B**) HEK293T cells were transfected with *BDNF* v11 5ʹ UTR-DLR wild-type and mutant constructs as indicated, and luciferase activity determined after 24 hours. For each mutant, the exon structure and number of functional uORFs (open circles) are indicated. (**C**) RT-qPCR was used to determine RLuc transcript levels normalized to FLuc expression. (**D**) Schematic of the 5ʹ UTR for the *BDNF* v14 transcript (NM_001143814.2). The pORF start codon is located in exon 3. (**E**) HEK293T cells were transfected with *BDNF* v14 5ʹ UTR-DLR wild-type and ΔExon2 constructs as indicated, and luciferase activity determined after 24 hours. Values are mean+SD, *n*=5 independent experiments for (**B**) and *n*=3 independent experiments for (**C** and **E**). Differences between groups were tested by Student’s *t*-test or one-way ANOVA and Bonferroni *post hoc* test, as appropriate. \**P*<0.05, \*\**P*<0.01, \*\*\**P*<0.001, \*\*\*\**P*<0.0001.

These data suggest that exon skipping strategies which aim to exclude *BDNF* v11 exon 2, or exons 2 and 3, from the mature *BDNF* transcript could potentially be exploited for therapeutic BDNF upregulation at the level of translation.

In contrast, analysis of *BDNF* v14 showed that when exon 2 was deleted (containing one uORF) there was no significant effect on reporter gene expression (**Figure 5D,E**). These data suggest that this transcript variant is not a suitable target for upstream exon skipping.

Based on these findings we selected *BDNF* v11 for further analysis. We hypothesised that the upregulation observed when exon 2 was deleted might be a consequence of the exclusion of 8 uORF sequences from the resulting ‘exon skipped’ mRNA. To this end, constructs were tested in which all 8 uORFs within exon 2 were disrupted by mutating their start codon ATG trinucleotides to TTG. Disruption of all eight uORFs located in exon 2 resulted in a ∼2.7-fold upregulation in reporter gene expression (**Figure 6A**), consistent with the relief of uORF-mediated repression. However, this upregulation was only a fraction (∼36%) of that observed when exon 2 was deleted in its entirety, suggesting that while uORFs contribute to pORF repression, additional regulatory elements may be contributing to its translational repression.

**Figure 6.**
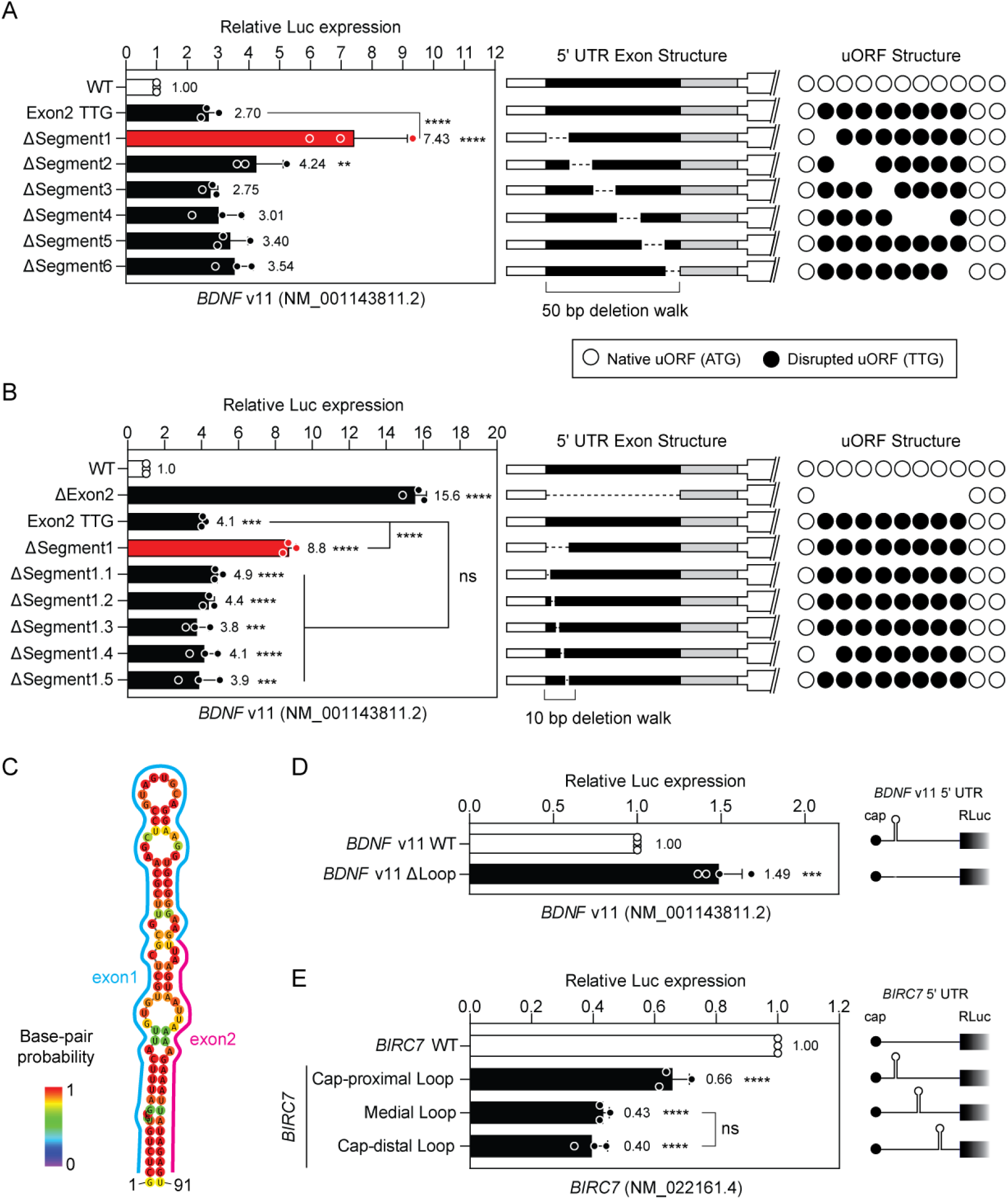
An RNA hairpin structure contributes regulated expression of *BDNF* v11. (**A**) HEK293T cells were transfected with *BDNF* v11 (NM_001143811.2) 5ʹ UTR-DLR wild-type and mutant constructs as indicated, and luciferase activity determined 24 hours post transfection. Constructs were generated in which 50 bp regions spanning exon 2 were sequentially deleted. (**B**) HEK293T cells were treated as above with *BDNF* v11 constructs in which 10 bp regions spanning the first 50 bp of exon 2 were sequentially deleted. (**C**) Structure of predicted RNA hairpin spanning the exon1-exon2 boundary. Dual luciferase assay data for plasmid constructs in which the putative hairpin is (**D**) deleted from the *BDNF* v11 5ʹ UTR, and (**E**) inserted into an unrelated 5ʹ UTR (of *BIRC7*, NM_022161.4) at positions as indicated. Values are mean+SD, *n*=3 independent experiments. Differences between groups were tested by Student’s *t*-test or one-way ANOVA and Bonferroni *post hoc* test, as appropriate. \*\**P*<0.01, \*\*\**P*<0.001, \*\*\*\**P*<0.0001, ns, not significant. Statistical comparisons are to the WT control group unless otherwise indicated.

We reasoned that other motifs located within might be responsible for the upregulation effect. To investigate this, a motif finding analysis was performed on the sequence of *BDNF* v11 exon 2 using BRIO (BEAM RNA Interaction mOtifs),(39) which identified two RNA binding motifs of interest (HuR #1 and HuR #2) (**Figure S1A**). However, deletion of these motifs, both individually and together, had no significant effect on pORF reporter expression (**Figure S1B**), suggesting that they are not responsible for the translation repressing activity contained within exon 2.

We next performed deletion walks to identify a nucleotide sequence in the *BDNF* v11 exon 2 that could account for its translation repressive activity. Mutant constructs were generated in which 50 bp contiguous regions of exon 2 were sequentially deleted. For the purpose of this experiment, all uORFs within exon 2 were disrupted, such that non-uORF repressive elements could be potentially identified. Deletion of the first 50 bp (ΔSegment 1) resulted in a pronounced 7.4-fold upregulation in reporter activity relative to the control construct where all the uORFs in exon 2 were disrupted (*P*<0.0001) (**Figure 6A**). These data suggest that the combination of uORF-mediated repression, together with a further regulatory element contained within these first 50 bp accounts for the majority of the repressive activity of this exon. A second series of deletion mutants were generated whereby 10 bp regions were deleted across the 50 bp region of interest at the start of exon 2. No upregulation activity was observed for any of these constructs (**Figure 6B**). Based on these data, we postulated that an RNA structural element, rather than a short sequence motif, might be responsible for the repressive activity mediated by the presence of *BDNF* v11 exon 2.

### Analysis of RNA structure in the *BDNF* v11 5ʹ untranslated region

RNA secondary structures were predicted for the *BDNF* v11 transcript using RNAfold,(40) together with all of the various deletion constructs. The WT *BDNF* v11 transcript was predicted to contain a prominent secondary structure motif that spanned the exon1-exon2 boundary (**Figure 6C**). This structure consisted of an imperfect 91 nucleotide hairpin with a stem length of 42 nucleotides and a minimum free energy of −23.5 kcal/mol. This structural motif was predicted to be absent when either exon 2 was deleted, or the first 50bp of exon 2 were deleted, but was broadly maintained in other constructs (**Figure S2**), suggesting that this element may be contributing to *BDNF* v11 translational repression.

Deletion of the hairpin resulted in a ∼1.5-fold de-repression of downstream reporter expression (*P*<0.001), consistent with it acting as a repressive element (**Figure 6D**). Notably, the magnitude of upregulation was less than that observed for the smaller deletions described above (**Figure 6A,B**), potentially because larger deletions may be impacting other regulatory signals within the *BDNF* v11 5ʹ UTR. To investigate the capacity of the *BDNF* v11 5ʹ UTR hairpin to induce translational repression in other contexts, we generated constructs in which the hairpin structure was inserted into the 5ʹ UTR of an unrelated gene. The *BIRC7* gene (NM_022161.4) was selected, as its 5ʹ UTR lacked any predicted uORFs. Insertion of the *BDNF* hairpin into the *BIRC7* 5ʹ UTR resulted in significant (*P*<0.001) repression of downstream reporter expression (**Figure 6E, Figure S3**). Repression was observed irrespective of whether the hairpin was inserted in the cap-proximal, medial, or cap-distal positions with respect to the 5ʹ cap, although repression was strongest in the medial and cap-distal positions (∼3-fold down-regulation). Taken together, these data suggest that a combination of an RNA secondary structure element, uORFs, and potentially other factors, combine to strongly repress the translational output of the *BDNF* v11 transcript.

### Characterisation of *BDNF* transcript expression in human brain and brain-derived cell cultures

We next aimed to determine whether the transcript isoforms investigated above are expressed in human brain via qualitative inspection of publicly-available RNA-Seq data from the developing human brain (covering transitions from embryonic, fetal, infancy, and adolescence stages)(41) and long-read RNA sequencing data from the brain of an adult Alzheimer’s disease patient. RNA-Seq read density was observed at 5ʹ UTR exons for RefSeq transcripts: NM_001709.5, NM_170731.5, NM_170733.4, NM_170734.4, NM_001143813.2, and NM_00143807.2 (also known as *BDNF* transcript variants v4 (MANE), v3, v5, v6, v13, and v9, respectively, **Figure 7A**). Detection of some of these transcript variants was experimentally confirmed in human brain RNA by reverse transcriptase-droplet digital PCR (RT-ddPCR), although the most highly abundant transcript isoform was found to be NM_170732 (v2) in contrast with the RNA-Seq data (**Figure 7B**).

**Figure 7.**
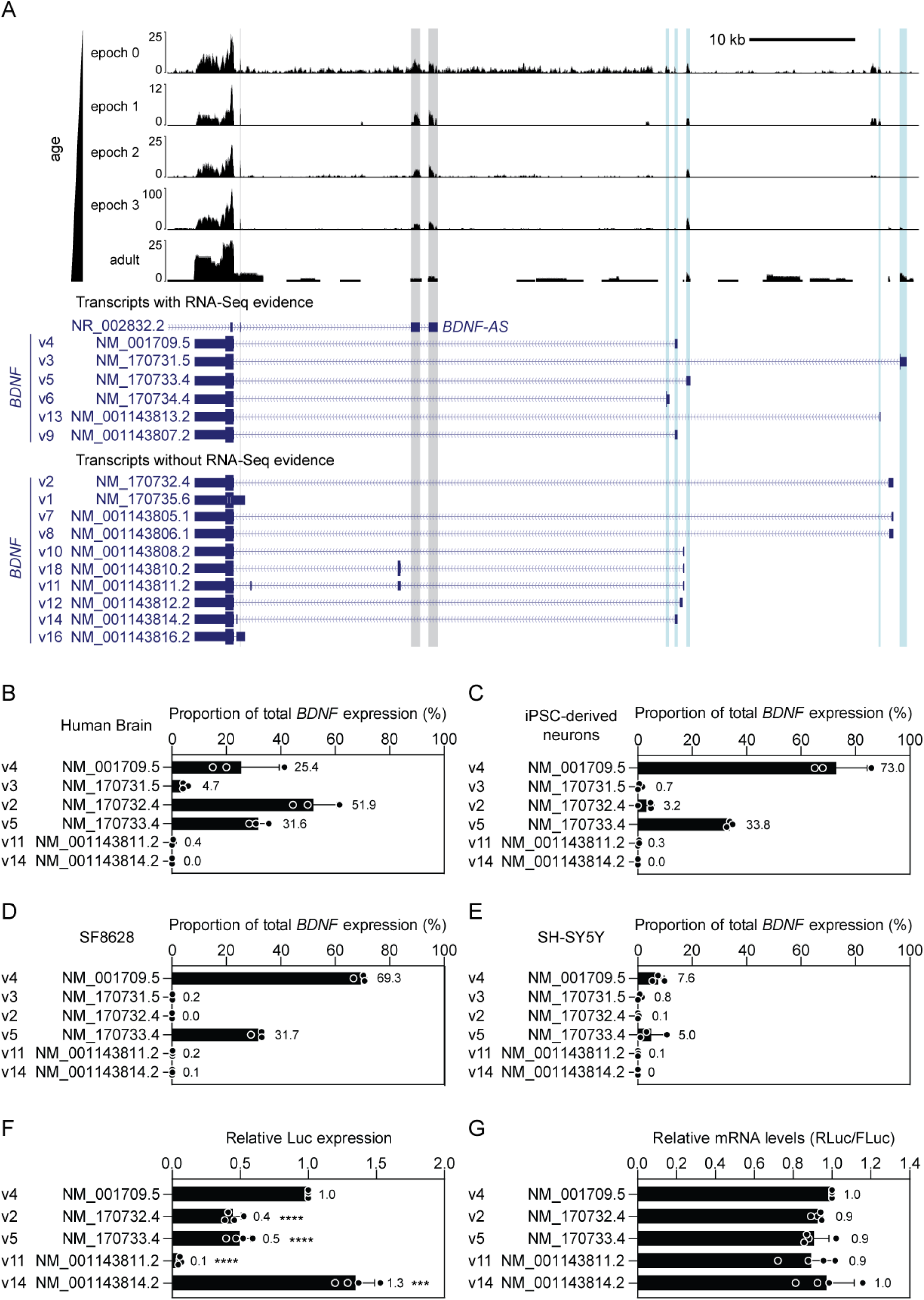
Characterisation of *BDNF* transcript isoform expression. (**A**) Publicly available RNA-Seq from the developing brain and long-read sequencing data from adult human brain were visualised for the *BDNF* locus. Annotated transcript isoforms were sorted according to whether there is empirical evidence for their expression in human brain. RNA-Seq data were aggregated from individual libraries (*N*=176) that were separated according to developmental epochs (0=embryonic/fetal [*n*=9], 1=fetal [*n*=83], 2=late fetal transition/infancy [*n*=39], 3=infancy/childhood/adolescence [*n*=45]). The adult sample is derived from an Alzheimer’s patient brain. Blue vertical bars highlight exons with evidence of expression. Grey bars highlight read density associated with an overlapping antisense long non-coding RNA gene (*BDNF-AS*). Expression of *BDNF* transcripts was determined by RT-ddPCR for (**B**) human brain (age 30 female), (**C**) iPSC-derived motor neuron cultures (*n*=3 lines), (**D**) SF8628 paediatric diffuse intrinsic pontine glioma cells, and (**E**) SH-SY5Y neuroblastoma cells. (**F**) HEK293T cells were transfected with dual luciferase reporter constructs carrying various wild-type *BDNF* promoter variants and luciferase activity assessed 24 hours later. (**G**) RNA was extracted from parallel cultures and the ratio of Renilla and firefly luciferase transcripts determined by RT-qPCR. Values are mean+SD. Differences between groups were tested by one-way ANOVA and Bonferroni *post hoc* test, \*\*\**P*<0.001, \*\*\*\**P*<0.0001. *n*=3 healthy cell lines for iPSC data. *n*=3 technical replicates for other cell lines and human brain RNA data. *n*=4 independent experiments for luciferase and RT-qPCR data.

The same RT-ddPCR assays were applied to a variety of human cell models including iPSC-derived motor neurons (**Figure 7C**), SF8628 (**Figure 7D**), and SH-SY5Y cells (**Figure 7E**). The most highly abundant transcript isoform was found to be the MANE variant (v4) in these cells, followed by NM_170733 (v5). Notably, the ‘skippable’ *BDNF* transcripts were expressed at very low, or undetectable levels, in both human brain RNA and total RNA from the various cell models, meaning that these transcripts are unlikely to be suitable targets for therapeutic BDNF upregulation.

We next compared the translational activity of the various wild-type 5ʹ UTRs previously investigated (**Figure 7F**) and compared these to the relative luciferase output from the MANE variant (NM_001709, v4). Interestingly, expression was reduced by ∼60% in variants NM_170732 (v2) and NM_170733 (v5) (*P*<0.0001), suggesting that while NM_170732 may be expressed at higher levels than the MANE variant in human brain, it exhibits a lower level of translational activity.

*BDNF* v11 (NM_001143811) was found to exhibit ∼90% less expression than the MANE variant (*P*<0.0001) (**Figure 7F**), consistent with our observations that this transcript has high potential for upregulation via upstream exon skipping (**Figure 5B**). However, the low levels of expression of this transcript preclude this as a therapeutic target in human brain (**Figure 7A-E**).

Interestingly, *BDNF* v14 (NM_001143814) was found to be ∼30% more translationally active than the MANE variant (*P*<0.001). This finding suggests that low abundance transcripts (such as NM_001143814) may nevertheless exhibit high translational activity. (**Figure 7F**). The skipping of exon 2 in *BDNF* v14 effectively results in the generation of the MANE variant *BDNF* v4, and so this result is consistent with the results shown in (**Figure 5E**). Luciferase transcript levels in cultures transfected with the various wild-type 5ʹ UTR constructs were not significantly changed (**Figure 7G**), suggesting that the differences in luciferase activity observed between 5ʹ UTR constructs occur at the level of translation.

### BDNF protein upregulation via base editing-mediated disruption of a uORF start codon

Based on the uORF validation data (**Figure 2B**) and experimentally observed transcript expression patterns in human brain and brain-derived cell models (**Figure 7**), we selected uORF-2 in *BDNF* v4 for further investigation. To maximise the chance of experimental success, we utilised HeLa cells to test experimental uORF manipulation approaches based on their relative ease of transfection. The *BDNF* v4 transcript was found to be the major expressed isoform in HeLa cells (**Figure 8A**), exhibiting a pattern of expression that was similar to that observed for iPSC-derived motor neurons (**Figure 7C**).

**Figure 8.**
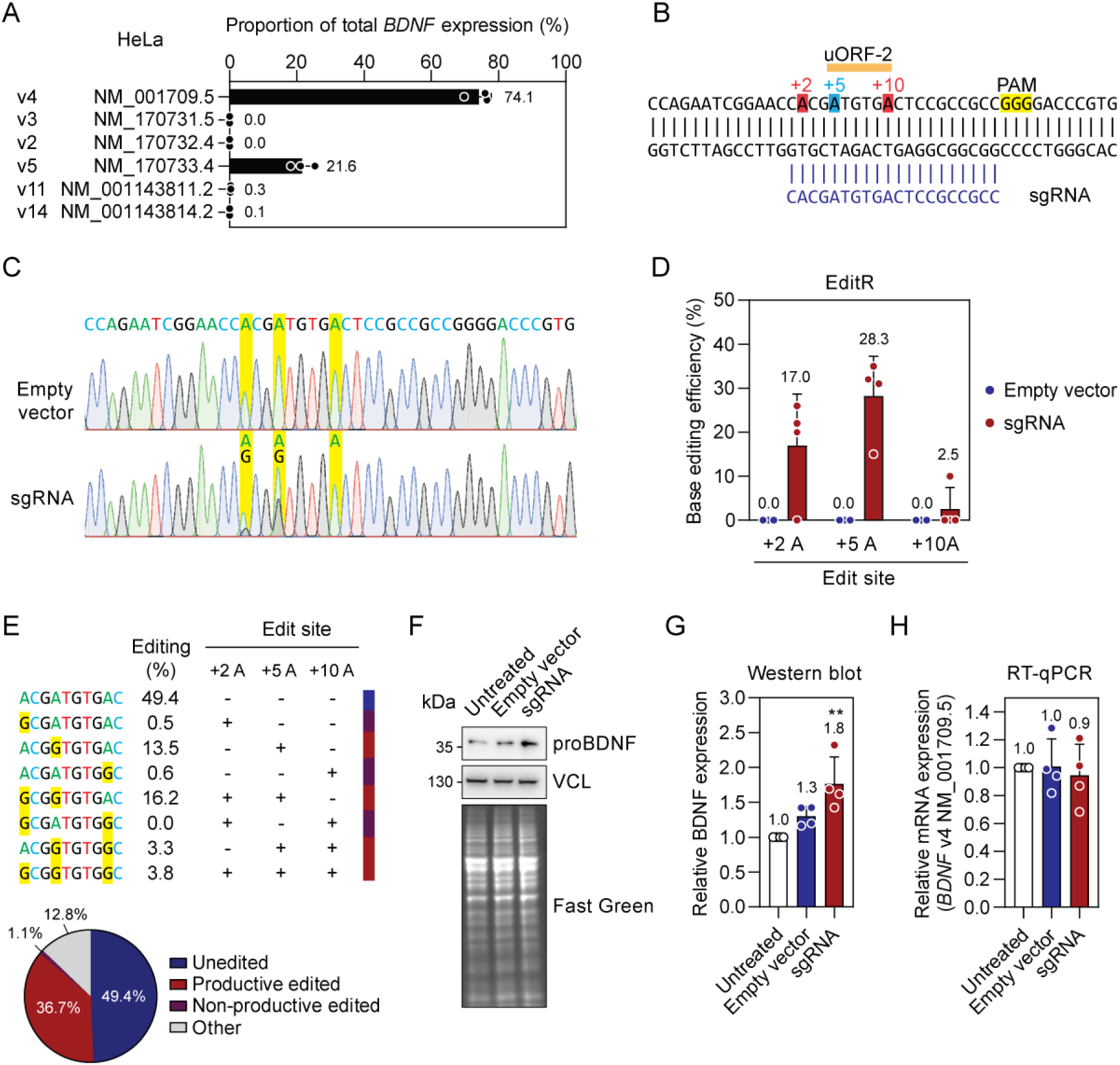
Upregulation of BDNF protein following base editing-mediated disruption of a uORF start codon. (**A**) Expression of *BDNF* transcript isoforms was determined by RT-ddPCR for HeLa cells. (**B**) Schematic of CRISPR-base editing strategy targeting the start codon of the *BDNF* v4 uORF-2. HeLa cells were transfected with plasmids encoding a nickase SpCas9 fused to an adenosine deaminase (ABE8e) and a single-guide RNA (sgRNA). Cells were dissociated three days post transfection, counted and re-seeded at a defined cell density. Materials was collected for further analysis after an additional 12 hours. (**C**) Representative Sanger sequencing chromatograms for control (Empty vector) and on-target sgRNA-treated cells. The target adenosine and potential bystander adenosines in the base editing window are highlighted in yellow. (**D**) Quantification of adenine base editing by EditR analysis. (**E**) Amplicon-seq analysis quantifying on-target and bystander editing events. (**F**) Representative western blot image showing BDNF protein detection. Equal protein loading was determined by blotting for vinculin (VCL) and by total protein staining using Fast Green. (**G**) Western blot quantification. (**H**) RT-qPCR analysis of *BDNF* transcript levels, normalized to *ACTB* expression. Values are mean+SD, *n*=4 independent experiments. Differences between groups were tested by one-way ANOVA and Bonferroni *post hoc* test, \*\**P*<0.01.

A CRISPR-nCas9 adenine base editing strategy using ABE8e(42) was devised, targeting *BDNF* v4 uORF-2 (**Figure 8B**). HeLa cells were transfected with plasmids encoding ABE8e and a single guide RNA (sgRNA) targeting uORF-2. Cells were harvested after three days and re-seeded at a consistent cell density across groups. In parallel, DNA editing was assessed by Sanger sequencing. After a further 12 hours, cells were harvested for protein and RNA analyses. Untreated cells, and cells treated with ABE8e and an empty sgRNA vector plasmids (empty vector) served as negative controls. Successful base editing was apparent from inspection of Sanger sequencing chromatograms, with overlaid A and G peaks observed at the target +5 site. Notably, some bystander editing was apparent at the +2 site, but not at the +10 site (**Figure 8C**). Quantification of the editing using EditR(43) revealed a mean target editing of 28.3% (with a maximum of 35%) (**Figure 8D**). Mean bystander editing was quantified as 17% at the +2 site and 2.5% at the +10 site. No base editing was observed in empty vector-treated cultures. The sample with the highest on-target base editing efficiency was selected for further analysis by next generation amplicon sequencing. This sample was selected in order to best capture the most diversity in editing outcomes. Quantification of editing outcomes revealed that productive editing outcomes (where the uORF-2 ATG was edited) represented 36.7% of all reads (**Figure 8E**), which closely matched the Sanger sequencing quantification (**Figure 8C**). Non-productive editing (whereby bystander editing occurred without uORF ATG editing) was observed in 1.1% of reads. Target uORF editing was associated with a ∼1.8-fold (*P*<0.01) increase in BDNF protein (pro form) expression by a maximum of 2.3-fold, consistent with partial uORF-disruption (**Figure 8F,G**). *BDNF* v4 mRNA levels were not significantly changed by the treatment, suggesting that increased proBDNF protein levels are due to changes at the translational level, consistent with uORF disruption (**Figure 8H**).

Clonal HeLa lines containing the desired *BDNF* v4 uORF-2 (ΔuORF2), and non-edited control clones (*n*=3), were obtained by limiting dilution. proBDNF protein was significantly (*P*<0.05) upregulated in the ΔuORF2 lines by 1.43-fold when lysates were collected 12 hours after seeding (**Figure 9A**) but not after 24 hours (**Figure 9B**). We reasoned that the effects of uORF disruption may be being overridden by another BDNF regulatory mechanism. To test this, wild-type HeLa cells were collected at 12, 24, 48, and 72 hours after seeding. proBDNF expression was significantly (*P*<0.05) upregulated by 2.2-fold at 72 hours (**Figure 9C,D**), whereas *BDNF* v4 transcript levels exhibited a reciprocal pattern of progressive downregulation (**Figure 9E**). These data suggested that cell crowding might have an effect on BDNF expression that is overwhelming the uORF regulation effect. To test this hypothesis, wild-type HeLa cells were seeded at four different densities (0.5, 1, 2, and 4×10^5^ cells/ml) and cells harvested either 12 or 24 hours later. In the case of both time points, a progressive increase in BDNF protein expression was observed (**Figure 9F,G**), while the opposite effect was observed for *BDNF* v4 transcript levels. Taken together, these data show that in cycling HeLa cells, *BDNF* expression is regulated in a cell density-dependent manner, and that the magnitude of this effect is sufficient to overwhelm the upregulation effect observed following targeted disruption of *BDNF* v4 uORF-2.

**Figure 9.**
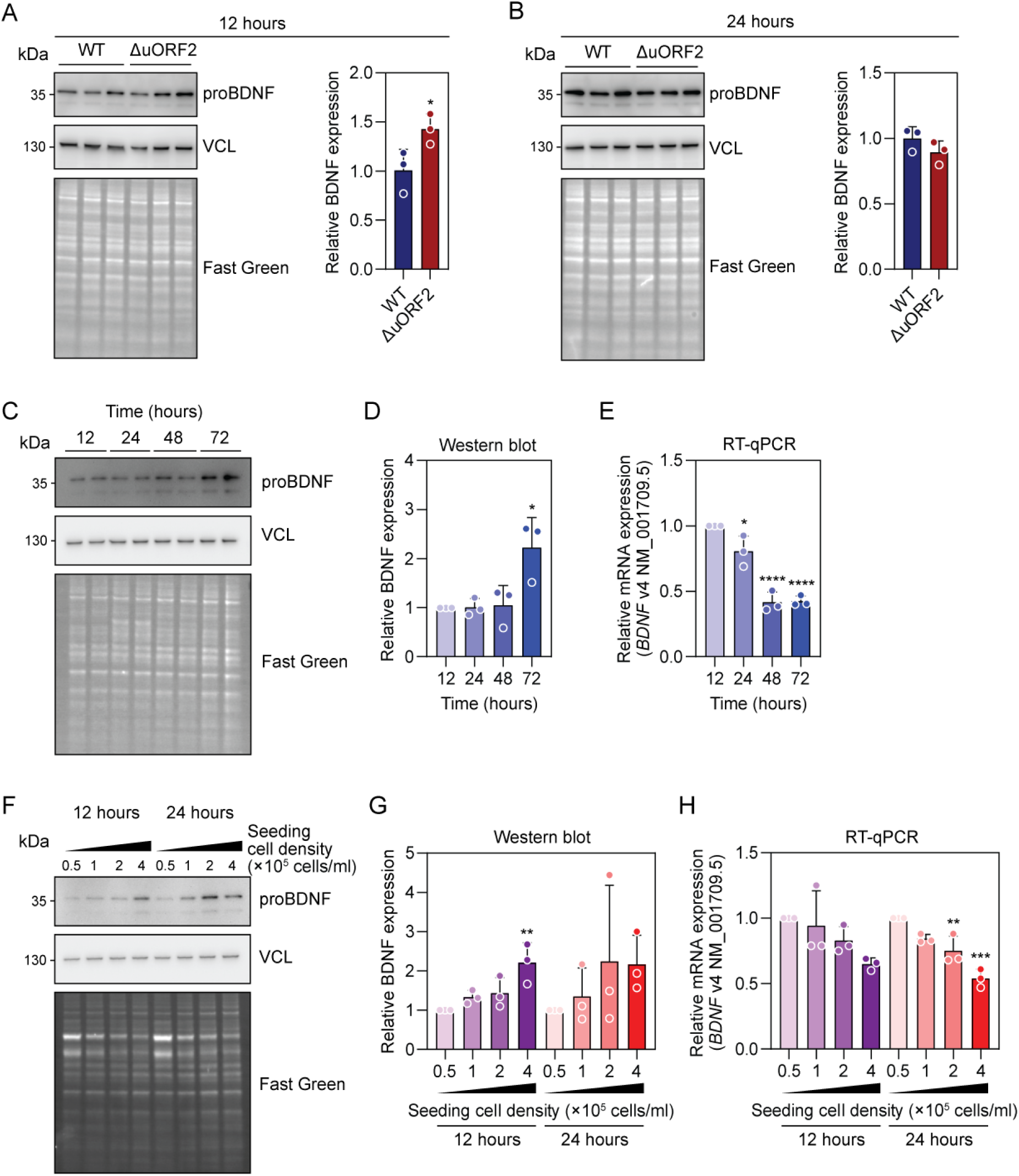
Investigation of uORF-disruption mediated BDNF upregulation in edited HeLa cell clones. Isolated HeLa cell clones carrying a uORF disrupting single base edit (ΔuORF2) were propagated, seeded, and the harvested at (**A**) 12 hours, or (**B**) 24 hours. BDNF protein expression determined by Western blot. Unedited clones containing the wild-type (WT) *BDNF* uORF served as controls. Wild-type HeLa cells were seeded and lysates collected 12, 24, 48, and 72 hours later. (**C**) BDNF expression was determined by western blot (with duplicate well technical replicates), and (**D**) quantified by densitometry. (**E**) *BDNF* v4 transcript levels were analysed by RT-qPCR in parallel. Wild-type HeLa cells were seeded at starting cell densities of 0.5, 1, 2, and 4 ×10^5^ cells/ml and lysates harvested 12 or 24 hours later. (**F**) BDNF expression was determined by western blot, and (**G**) quantified by densitometry. (**H**) *BDNF* v4 transcript levels were analysed by RT-qPCR in parallel. Values are mean+SD. For (**A**-**B**), differences between groups were assessed by one-tailed Student’s t-test, *n*=3 separate clones, each reported as the mean of *n*=2-3 independent experiments. For (**C-H**), differences between groups were tested by one-way ANOVA and Bonferroni *post hoc* test. *n*=3 independent experiments, \**P*<0.05, \*\**P*<0.01, \*\*\**P*<0.001, \*\*\*\**P*<0.0001.

## Discussion

The *BDNF* gene exhibits a complex transcriptional landscape with the potential to generate multiple transcript isoforms. The majority (14 out of 17 RefSeq annotated transcripts) are predicted to contain at least one uORF (**Figure 1**, **Table 1**). Five *BDNF* transcripts were selected for experimental characterisation, all of which were validated as being uORF-regulated to some extent, suggesting that uORF regulation plays an important role in shaping the translational output of the *BDNF* locus (**Figures 2-5**). However, the pattern of uORF regulation differed between transcripts, and a number of interesting features were observed. For transcripts containing multiple uORFs, we typically observed additive effects of having multiple uORFs (**Figure 2,3**). We also observed redundancy between uORF activity (**Figure 3,4**), and that not all uORFs are equal, with some exhibiting a greater contribution to pORF repression than others (**Figure 2,3**). Moreover, we observed that the minimal uORF (i.e. methionine-stop) is capable of functioning as a repressive uORF (**Figure 2**).

The multitude of uORF-regulated transcripts originating from the *BDNF* locus present exciting opportunities for therapeutic manipulation. It has previously been reported by Liang *et al*., that uORF activity can be disrupted to activate pORF translation using steric block antisense oligonucleotides (ASOs) targeted to uORF start codons.(44) There have been since been conflicting reports regarding the capability of ASOs to interfere with uORF function. We have been unable to reproduce some of the main findings from this study by targeting the *RNASEH1* uORF using the same ASO sequences reported by the authors.(45) Nevertheless, some studies have reported similar effects targeting 5ʹ UTRs with steric block antisense oligonucleotides,(46–48) However, some studies have reported uORF-independent protein upregulation effects that with ASOs overlapping uORFs,(49) and even pORF downregulation after direct uORF start codon targeting.(50) We have therefore been motivated to explore alternative approaches for therapeutic uORF interference.

Upstream exon skipping is a particularly attractive approach for counteracting the repressive effects of uORFs. There are multiple FDA-approved splice switching oligonucleotides (in non-uORF contexts), suggesting that this modality has reached a point of maturity.(51) Most notably, nusinersen (Spinraza) is a highly-effective splice switching oligonucleotide for the treatment of spinal muscular atrophy, which is an 18mer comprised of full phosphorothioate-2ʹ-*O*-methoxyethyl chemistry that is administered by the intrathecal route. The example of nusinersen has shown that this pattern of oligonucleotide chemistry and route-of-administration are viable approaches for the long-term treatment of infants and children. The widespread distribution of nusinersen throughout the CNS is highly encouraging for the treatment of brain indications, as would be required for *BDNF*-targeted therapies.(52) As such, nusinersen has served as a ‘template’ for other therapies. Most notably, milasen was developed as a personalised splice switching therapy for a single patient with CLN7 Batten disease modelled on nusinersen.(53) Similarly, the nusinersen template might easily be re-deployed to skip upstream 5ʹ UTR exons with the aim of disrupting uORF function. To this end, we explored two *BDNF* variants with ‘skippable’ upstream exons (**Figure 5**). Deletion of exon 2 of *BDNF* v11 resulted in a profound upregulation effect, thereby validating this approach. However, inspection of cortex RNA-Seq data from developing cortex, together with absolute quantification ddPCR in adult human brain and brain-derived cell lines suggests that this transcript is present at very low levels or not detected (**Figure 7**). As such, splice manipulation of low levels of this mRNA are unlikely to have a significant effect on bulk BDNF protein output. However, such an upstream exon skipping approach is likely to be applicable in other genetic/therapeutic contexts. Notably, a similar effect has been reported by Ang *et al*.(54) Additionally, we found that other factors, such as the presence of RNA structural motifs, might also be disrupted through upstream exon skipping, expanding the possible utility of this approach.

By contrast, deletion of exon 2 in *BDNF* v14 resulted in a slight decrease in reporter expression. These findings demonstrate that while the removal of one of more exons has the potential to exclude negative regulatory information from the mature transcript, it might equally remove positive regulatory information, leading to a decrease in target protein expression. Interestingly, skipping of *BDNF* v14 exon 2 effectively converts this transcript isoform into another annotated isoform, *BDNF* v4. We have demonstrated that this transcript contains a potent uORF in exon 1, and that skipping of exon 2 decreases the intercistronic distance between the uORF and pORF, which may account for the observed increase in pORF repression.(55, 56) These findings, which are not immediately obvious, suggest that uORF therapeutic targets must be functionally validated to confirm the intended direction of effect. Furthermore, our data highlights the importance of validating the presence of the target transcript in disease-relevant tissues, as considerable variability in exon inclusion and transcription start site usage is often observed at gene 5ʹ ends.

We next sought to utilise a base editing strategy for targeted uORF manipulation. Transient transfection of HeLa cells with the ABE8e adenine base editor and a uORF-targeting guide RNA resulted in up to 35% editing and a corresponding ∼1.8-fold activation of BDNF protein expression in transiently transfected cells (**Figure 8**) and ∼1.4-fold activation in clonally isolated cultures. These data provide proof-of-concept evidence that uORFs can be manipulated for therapeutic purposes in the specific case of *BDNF* (**Figure 9**). Interestingly, when cells were cultured for prolonged periods the effects of uORF-disruption-mediated BDNF upregulation were lost, which we attribute to a cell density-associated upregulation of BDNF expression which overwhelmed the uORF effect (**Figure 9**). This aspect of BDNF regulation must be carefully considered when advancing this approach in more physiologically-relevant cell models and small animal studies.

Importantly, editing takes place in non-protein-coding regions, where off-target effects are expected to have limited impact. An exciting possibility is that such an approach would have broader applicability, especially considering the abundance of predicted uORFs in the human transcriptome. Indeed, CRISPR-Cas9-mediated uORF disruption has been reported in plants,(57, 58) and in human cells.(59) An obvious extension of the approach described here would be to use RNA editing technologies for transient targeting of a *BDNF* uORF. A variety of oligonucleotide triggers have been proposed with various designs and chemical compositions.(60–62) Such a technology would be in many ways preferable, as editing trigger oligonucleotides could be introduced via intrathecal injection, with the degree of target protein upregulation tuneable and/or reversible through the control of dosing. Similarly, expressed RNA editing triggers have also been described,(63) which may have similar upregulation potential if deployed appropriately.

Further therapeutic translation of *BDNF* targeting strategies will likely necessitate the use of rodent models, and thereby modification of the effector molecules will be needed so that they can target the corresponding mouse or rat sequences. Alternatively, small regions of the rodent 5ʹ UTR could be humanised in order to enable further translational development.

In summary, this study identified *BDNF* as a promising target for uORF manipulation therapies. We report multiple new regulatory features of *BDNF* transcript regulation and describe two possible approaches for achieving therapeutic target upregulation. Using base editing we were able to demonstrate robust BDNF protein activation even after transient transfection and ∼28% editing at the target uORF start codon adenosine. These studies provide a template for the application of these technologies for the upregulation of other therapeutic target genes.

## Methods

### Bioinformatics and public datasets

uORFs were predicted using custom in-house software. Ribo-Seq data and RNA-Seq data were obtained from GWIPS-Viz.(64, 65) Visualisation of Ribo-Seq/RNA-Seq data in RNA space was performed using custom in-house software. Details of *BDNF* transcript isoforms were downloaded using the Table Browser function of the UCSC Genome Browser.(66) RNA-Seq data from the BrainVar consortium was used to determine *BDNF* transcript isoform expression.(41) These data are rRNA-depleted (by ribozero) bulk tissue libraries derived from human dorsolateral prefrontal cortex (*N*=176). Data were stratified by developmental stage (i.e. epoch). Epoch 0: first trimester, epoch 1: second trimester, epoch 2: third trimester and infancy, and epoch 3: infancy to adult. Long read RNA sequencing data (PacBio platform) from an adult brain from an Alzheimer’s disease patient were obtained from the PacBio website (https://downloads.pacbcloud.com/public/dataset/Alzheimer2019_IsoSeq/). Motif finding analysis was performed using BRIO.(39) RNA structure analysis and visualisation was performed using RNAfold.(40)

### Human Brain RNA

Human brain RNA was obtained from Thermo Fisher Scientific (product no: AM7962, lot no: 2661603). The sample was obtained from a 30-year-old Caucasian female (cause of death: anoxia/asphyxiation).

### Reporter Plasmid Generation

Plasmid constructs were generated in-house by Gibson assembly or generated as a service by Azenta Life Sciences (Manchester, UK). A custom generated dual luciferase reporter was used for uORF validation. This vector consists of two independent reporter transgene cassettes; firefly luciferase (FLuc) and Renilla luciferase (RLuc).

### Cell Culture

All immortalised cells (HEK293T, SH-SY5Y, SF8628, and HeLa) were cultured in a humidified incubator at 37°C (5% CO_2_) in Dulbecco’s Modified Eagle Medium (DMEM-Glutamax) that was supplemented with 10% Fetal Bovine Serum (FBS) (both Thermo Fisher Scientific) and 1% Penicillin, Streptomycin and Amphotericin B (PSA) (Merck Life Science, Gillingham, UK). Cell cultures were confirmed free of mycoplasma contamination through monthly testing.

For reporter construct assays, cells were transfected with plasmids using Lipofectamine 2000 (Thermo Fisher Scientific) or polyethylenimine (PEI). For Lipofectamine 2000 transfection, plasmids and transfection reagent were prepared separately in Opti-MEM, before being combined, incubated for 10 minutes, and then added to cells in a dropwise manner, as according to manufacturer’s instructions. A ratio of 1 µl of Lipofectamine 2000 to 1 µg of plasmid DNA was used. For PEI transections, a weight:weight ratio of 4:1 transfection reagent:plasmid DNA was used. PEI and plasmid DNA was prepared separately in DMEM, combined, and then incubated for 20 minutes, before dropwise addition to cells. A total of 100 ng or 500 ng plasmid DNA was used for 96-well and 24-well plates, respectively.

For base editing experiments, transfection was performed using Lipofectamine 2000 with a ratio of 2.1 µl transfection reagent per 1 µg of plasmid DNA. A total of 2.5 µg of plasmid was transfected per well of a 24-well plate.

### iPS Cells

The iPSC lines used in this study were derived from skin biopsy fibroblasts, collected under ethical approval granted by the South Wales Research Ethics Committee (WA/12/0186) in the James and Lillian Martin School Centre for Stem Cell Research, University of Oxford, under standardized protocols which we have described elsewhere.(67) Fibroblasts and derived iPSC lines tested negative for mycoplasma (MycoAlert, Lonza, UK). The iPS cells were differentiated into motor neurons *in vitro* according to our previously published methods.(67) Briefly, the iPS cells were cultured on Geltrex in mTESR 1 supplemented with mTESR supplement (both Stem Cell Technologies, Cambridge, UK) and antibiotics (Thermo Fisher Scientific). Induction media was added to the iPS cells (at 90% confluence) and consisted of DMEM/F12/Neurobasal medium 1:1, 1× N2 supplement, 1× B27 supplement (all Thermo Fisher Scientific), ascorbic acid (0.5 μM, Sigma-Aldrich, Gillingham, UK), β-mercaptoethanol (50 μM, Thermo Fisher Scientific), Compound C (1 μM, Bio-Techne, Abingdon, UK), Chirr99201 (3 μM, Bio-Techne). On the second day, the induction medium was supplemented with all-*trans* retinoic acid (1 μM, Sigma-Aldrich) and Smoothened Agonist (500 nM, Bio-Techne). Two days later, Chirr99201 and Compound C were removed from the medium. On the 9^th^ day of differentiation, neural progenitor cells were split 1:3 using Accutase (Thermo Fisher Scientific) and ROCK inhibitor (Bio-Techne) was added for 24 hours. On day 19, the progenitors were split in their final plating conditions and the medium was supplemented with BDNF (10 μM, Thermo Fisher Scientific), GDNF (10 μM, Thermo Fisher Scientific), and laminin (500 ng/µl). DAPT (10 μM, Bio-Techne) and ROCK inhibitor were added for the first 7 days, and then removed for the remainder of the maturation. RNA was collected between days 30 and 35.

### Dual Luciferase Assay

Luciferase activity was determined using the Dual-Glo Luciferase assay (Promega) according to manufacturer’s instructions. Briefly, HEK293T cells were transfected with plasmids as appropriate in quadruplicate. 24 hours after transfection, half of the media volume was removed, and Dual-Glo Reagent was added to each well. Samples were incubated at room temperature for 10 minutes and then the cell lysates transferred to a flat-bottomed Greiner 96-well plate (Merck). FLuc activity was measured, followed by quenching with an equal volume of Stop & Glo Reagent, a second 10-minute incubation at room temperature, and then measurement of RLuc activity. Luciferase activities were recorded on a CLARIOstar luminometer (BMG Labtech, Aylesbury, UK) using a focal height of 9.0 mm and the no filter setting. The Renilla luciferase activity was normalized to the firefly luciferase activity for each sample, and the data were scaled such that the mean value of the control group was returned to a value of 1.

### RT-qPCR

To quantify firefly and Renilla transcripts, total RNA was first extracted using the Maxwell RSC Instrument and the Maxwell RSC simplyRNA Tissue Kit (both Promega). Reverse transcription was performed using the High-capacity cDNA reverse transcription kit (Thermo Fisher Scientific) as according to manufacturer’s instructions using 200-1000 ng (typically 400 ng) input total RNA.

cDNA was diluted 1:5 in nuclease-free water prior to quantitative polymerase chain reaction (qPCR) amplification using Power SYBR Green PCR Master Mix and the StepOnePlus Real-Time PCR System (both Applied Biosystems, Warrington, UK). Universal cycling conditions were used (i.e. 95°C for 10 minutes, followed by 40 cycles of 95°C for 15 seconds and 60°C for 1 minute) and samples were run in duplicate. 10 pmols of each primer were included per reaction. Reaction specificity was confirmed by post-run melting curve analysis. Relative quantification was performed using the Pfaffl method.(68) Primer sequences are listed in **Table S1**.

### RT-ddPCR

Absolute quantification of RNA transcript isoforms was determined by reverse transcription-droplet digital PCR (RT-ddPCR), using diluted cDNA (1:5, prepared as described above). Reactions were prepared using QX200 ddPCR EvaGreen Supermix (Bio-Rad) and 10 pmols of each primer per 20 µl reaction. Droplets were prepared using the QX200 Droplet Generator and QX200 Droplet Generation Oil for EvaGreen (both Bio-Rad). Subsequently, droplets were transferred to a 96-well plate and the plate sealed using the PX1 PCR Plate Sealer (Bio-Rad). PCR was performed using the following cycling conditions: 95°C enzyme activation step for 5 minutes followed by 40 cycles of a two-step cycling protocol (95 °C for 30 seconds and 60 °C for 1 minute). The ramp rate between these steps was set to 2°C per second. After thermal cycling, the plate was analysed using a QX600 droplet reader (Bio-Rad) according to manufacturer’s instructions.

### Base Editing

Base editing guide RNA sequences were designed manually by inspection of the sequence and SpCas9 PAM sequences (NGG) identified. The nSpCas9-ABE expression cassette was derived from the ABE8e plasmid(42) (Addgene, #138489) and cloned into a custom expression vector. Guide RNAs were cloned into a custom U6 promoter expression vector.

To assess base editing efficiency, genomic DNA was extracted using the Maxwell RSC Cell DNA Purification Kit (Promega) following manufacturer’s instructions. A region of the *BDNF* locus encompassing the edit site was amplified by PCR using Q5 High-Fidelity DNA Polymerase (New England Biolabs) and subject to Sanger sequencing (Azenta Life Sciences). Primer sequences for editing analysis are listed in **Table S1**. The proportion of edits was quantified using EditR.(43)

Next generation amplicon sequencing was performed using the Amplicon-EZ service (Azenta Life Sciences). Sequencing reads were processed using custom python scripts that used string matching to identify edits occurring at the target uORF site together with bystander edits. Reads that were not matched using these criteria were classified as ‘other’, and were either of atypical length or contained additional mismatches in non-target regions of the sequence.

### Western Blot

Cells were washed with PBS, and then lysed in RIPA buffer (Thermo Fisher Scientific) with cOmplete EDTA-free protease inhibitor (Roche, Welwyn Garden City, UK). The lysates were cleared by centrifugation at 12,000 *g* for 10 minutes at 4°C. Protein concentrations were determined by BCA Protein Assay (Thermo Fisher Scientific) according to manufacturer’s instructions. Equal amounts of total protein were loaded onto precast 10% NuPAGE Bis-Tris mini or midi gels (Thermo Fisher Scientific) and separated by sodium dodecyl sulfate–polyacrylamide gel electrophoresis (SDS-PAGE). Protein samples were electrotransferred onto a 0.2 µm PVDF (polyvinylidene fluoride) membrane (Merck Millipore, Watford, UK). Membranes were stained with Fast Green FCF (Sigma-Aldrich) and total protein imaged using the ChemiDoc MP Imaging System (Bio-Rad). Membranes were subsequently blocked by 5% milk in Tris-buffered saline supplemented with Tween 20 (TBST). Blocked membranes were incubated with primary antibodies overnight in 2% bovine serum albumin (BSA) in TBST at 4°C. The next day, membranes were washed three times with TBST (10 minutes each), followed by incubation with secondary antibodies for 1 hour at room temperature. Membranes were washed three times and signal developed using Clarity Western ECL Substrate (Bio-Rad). Blots were visualized using the ChemiDoc MP Imaging System. Details of antibodies are provided in **Table S2**.

### Statistical Analysis

Statistical analyses were performed using GraphPad Prism Software Version 10.1.2 (GraphPad Software Inc., San Diego, California, USA). For comparisons of two samples, a Student’s *t*-test was used. For comparisons of more than two groups, an ordinary one-way analysis of variance (ANOVA) was performed with Bonferroni’s *post hoc* test for inter-group comparisons.

## Acknowledgements

This work was supported by grants from Great Ormond Street Hospital Sparks Fund/Dravet Syndrome UK (awarded to MJAW and TCR), the Oxford University Press John Fell Fund and Medical Life Sciences Translational Fund (awarded to TCR).

## Author Contributions

TCR, NS, BH, DG, and MJAW conceived the study. TCR, DG, and MJAW supervised the work. NF, TG, NS, DY, and BH performed experimentation. AL and SJS provided informatics analysis. RD and KT provided iPSC material. TCR wrote the first draft of the manuscript. All authors contributed to the final version of the manuscript.

## Data Availability Statement

All data are included in the manuscript. Raw data are available on request.

