## Supplementary Material for "Targeted BDNF upregulation via upstream open reading frame disruption"

| Target | Forward | Reverse |
|---|---|---|
| **BDNF_total** | GGATGAGGACCAGAAAGTTCG | GGACATGTTTGCAGCATCTAGG |
| **BDNF_NM_001709** | GTGTGGACCCCGAGTTCC | CAGCCTTCATGCAACCAAAG |
| **BDNF_NM_170731** | GGGAGACGAGATTTTAAGACACT | CAGCCTTCATGCAACCAAAG |
| **BDNF_NM_170732** | CTGGGTTAACTTTGGGAAATGC | CAGCCTTCATGCAACCAAAG |
| **BDNF_NM_170733** | GCTGCCTTGATGGTTACTTTG | CAGCCTTCATGCAACCAAAG |
| **BDNF_NM_001143811** | TGCATTCTGACCTATTGACTGG | CAGCCTTCATGCAACCAAAG |
| **BDNF_NM_001143814** | GGACCCGTGAGGTTTGTG | GGTGGAACTGAGATTAGATGGC |
| ***ACTB*** | CACCATTGGCAATGAGCGGTTC | AGGTCTTTGCGGATGTCCACGT |
| **RLuc** | GTAACGCTGCCTCCAGCTAC | CCAAGCGGTGAGGTACTTGT |
| **FLuc** | ACTCTAAGACCGACTACCAGG | GTAGACCCAGAGCTGTTCATG |
| ***BDNF editing region*** | AAGCTCAACCGAAGAGCTAAA | AACTCCAAATCGTCCCTTCTAC |

**Table S1**

**RT-qPCR/RT-ddPCR assays and editing PCR primers used in this study.**

All sequences are written 5′ to 3′.

| Target | Host (clone) | Product ID | Manufacturer | Dilution |
|---|---|---|---|---|
| **Primary Antibodies** | | | | |
| **anti-BDNF [EPR1292]** | rabbit mAb | ab108319 | Abcam | 1:5,000 |
| **anti-VCL [hVIN-1]** | mouse mAb | V9131 | Sigma-Aldrich | 1:5,000 |
| **Secondary Antibodies** | | | | |
| **anti-rabbit IgG-HRP** | goat | 7074 | Cell Signalling | 1:2,000 |
| **anti-mouse IgG-HRP** | horse | 7076 | Cell Signalling | 1:2,000 |

**Table 2**

**Antibodies used in this study.**

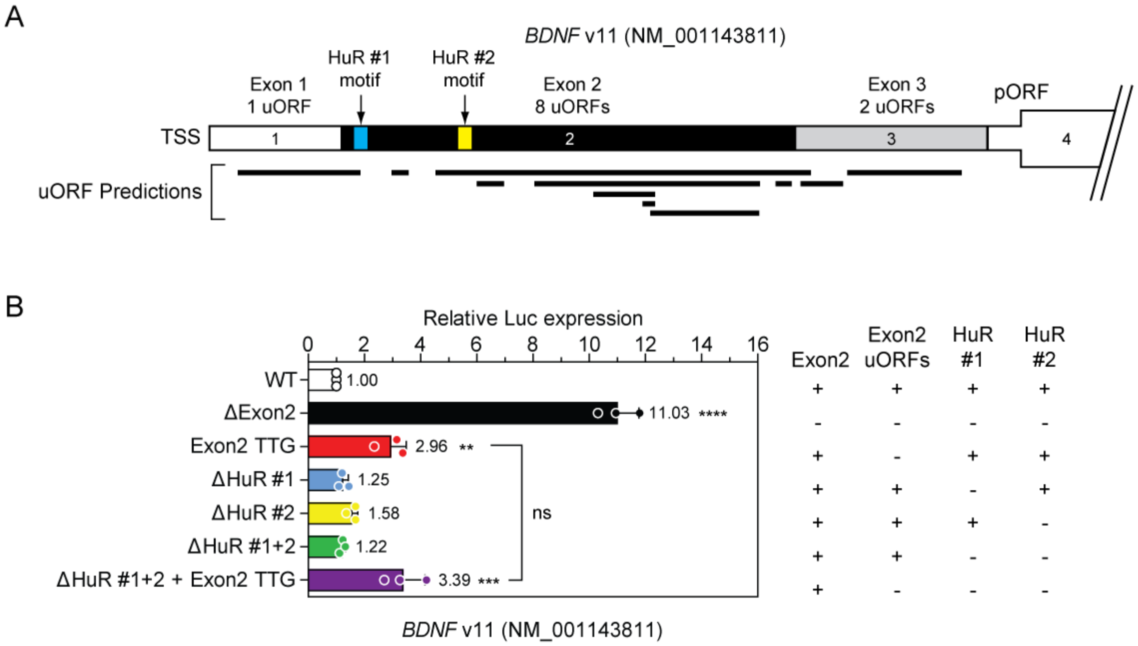


**Figure S1**

**uORFs are only partially responsible for the repressive activity of *BDNF* v11 exon 2.**

(**A**) Schematic of the *BDNF* v11 5′ UTR with the HuR #1 and HuR #2 motif sites indicated. The sizes and positions of exons, the locations of predicted uORFs, and the number of uORFs per exon are also indicated. (**B**) HEK293T cells were transfected with various *BDNF* v11 5′ UTR-DLR constructs. Mutants were generated in which either or both of the HuR motifs were deleted. An additional construct in which both motifs were deleted and all exon 2 uORFs were disrupted was tested in parallel. Luciferase activity was determined 24 hours post transfection. Values are mean+SD, $n$=3 independent experiments. Differences between groups were tested by one-way ANOVA and Bonferroni *post hoc* test. **$P$<0.01, ***$P$<0.001, ****$P$<0.0001, ns, not significant.

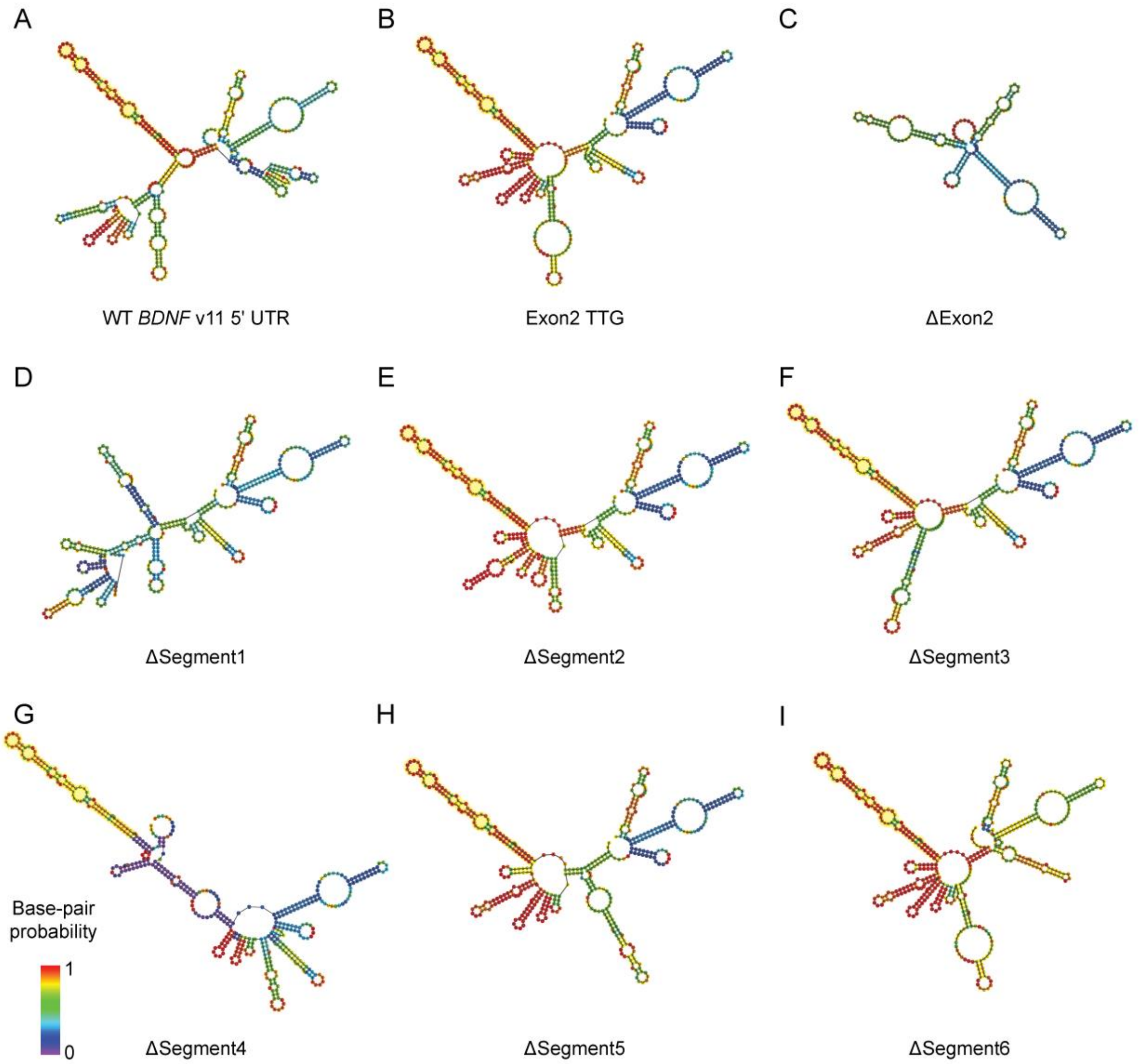


**Figure S2**

**RNA structure predictions for *BDNF* v11 and associated deletion constructs.**

RNAfold structures for (**A**) the wild-type *BDNF* v11 UTR, and the *BDNF* v11 5′ UTR whereby; (**B**) all eight exon2 uORFs are disrupted by ATG-to-TTG mutations, (**C**) Exon2 is deleted in its entirety, and (**D-I**) 50 nucleotide segments of exon2 are sequentially deleted (in the context of all exon2 uORFs being disrupted). The hairpin structure of interest is highlighted in yellow.

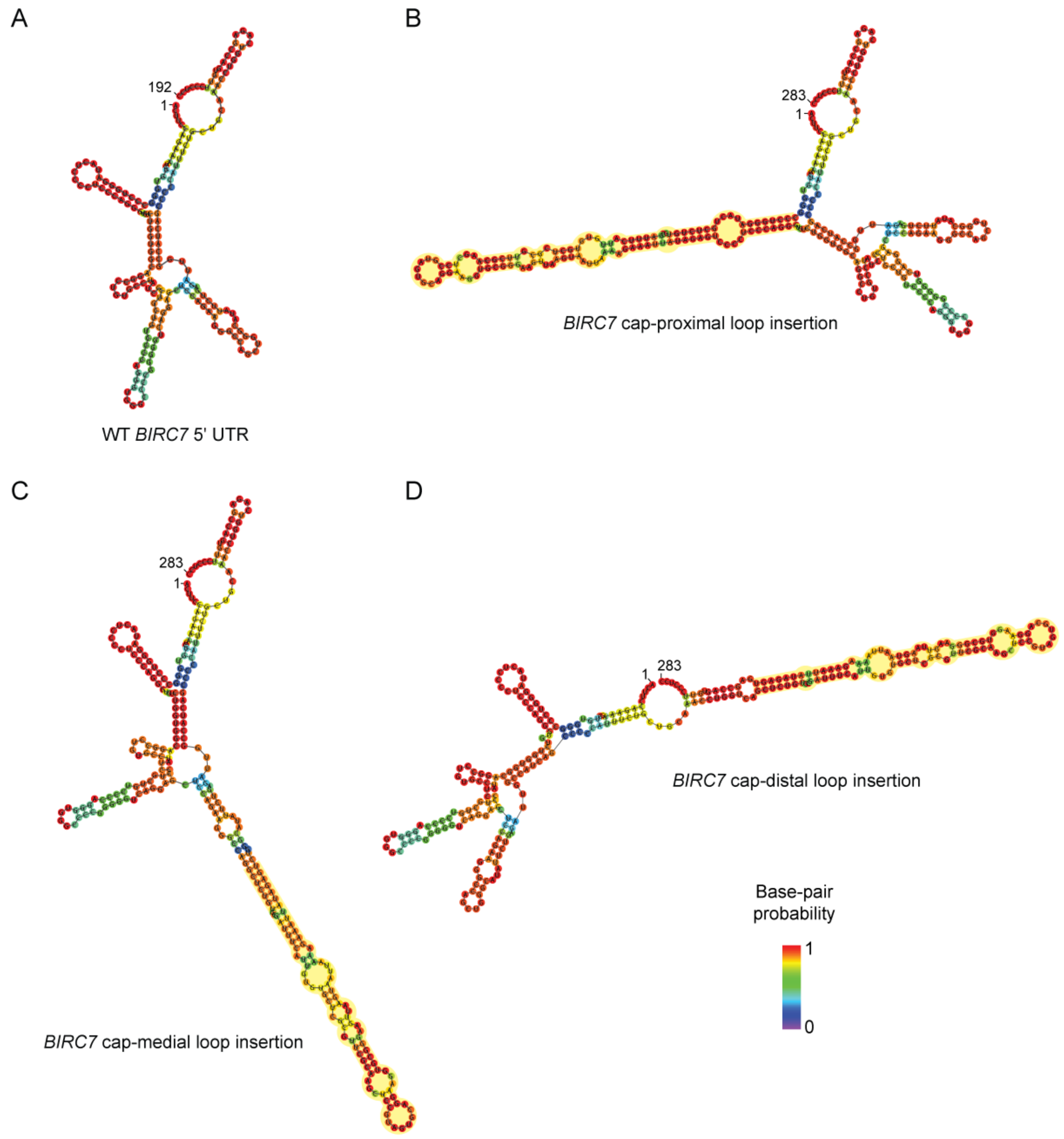


**Figure S3**

**RNA structure predictions for *BIRC7* 5′ UTRs with *BDNF* v11 loop insertions.**

RNAfold structures for (**A**) the wild-type *BIRC7* 5′ UTR, and constructs in which a *BDNF* v11-derived loop sequence (highlighted in yellow) is inserted in the (**B**) cap-proximal, (**C**) medial, and (**D**) cap-distal positions.
